# SARS-CoV-2 B.1.351 (beta) variant shows enhanced infectivity in K18-hACE2 transgenic mice and expanded tropism to wildtype mice compared to B.1 variant

**DOI:** 10.1101/2021.08.03.454861

**Authors:** Ferran Tarrés-Freixas, Benjamin Trinité, Anna Pons-Grífols, Miguel Romero-Durana, Eva Riveira-Muñoz, Carlos Ávila-Nieto, Mónica Pérez, Edurne Garcia-Vidal, Daniel Pérez-Zsolt, Jordana Muñoz-Basagoiti, Dàlia Raïch-Regué, Nuria Izquierdo-Useros, Ignacio Blanco, Marc Noguera-Julián, Victor Guallar, Rosalba Lepore, Alfonso Valencia, Júlia Vergara-Alert, Bonaventura Clotet, Ester Ballana, Jorge Carrillo, Joaquim Segalés, Julià Blanco

**Affiliations:** IrsiCaixa AIDS Research Institute, Germans Trias i Pujol Research Institute (IGTP), Can Ruti Campus, UAB, 08916, Badalona, Catalonia, Spain; Barcelona Supercomputing Center, 08034, Barcelona, Catalonia, Spain; IRTA, Centre de Recerca en Sanitat Animal (CReSA, IRTA-UAB), Campus de la UAB, 08193 Bellaterra, Spain; Germans Trias i Pujol Hospital, 08916, Badalona, Catalonia, Spain; Catalan Institution for Research and Advanced Studies (ICREA), 08010, Barcelona, Catalonia, Spain; University of Vic–Central University of Catalonia (UVic-UCC), 08500, Vic, Catalonia, Spain; Departament de Sanitat i Anatomia Animals, Facultat de Veterinària, UAB, 08193, Bellaterra, Catalonia, Spain

**Author notes:** **Corresponding author:** Julià Blanco, PhD, Senior Researcher, Institut de Recerca de la Sida. IrsiCaixa, IGTP, Hospital Germans Trias i Pujol, Ctra. de Canyet s/n. 2a Planta Maternal. 08916 Badalona. Barcelona.

## Abstract

SARS-CoV-2 variants display enhanced transmissibility and/or immune evasion and can be generated in humans or animals, like minks, thus generating new reservoirs. The continuous surveillance of animal susceptibility to new variants is necessary to predict pandemic evolution. In this study we demonstrate that, compared to the B.1 SARS-CoV-2 variant, K18-hACE2 transgenic mice challenged with the B.1.351 variant displayed a faster progression of infection. Furthermore, we also report that B.1.351 can establish infection in wildtype mice, while B.1 cannot. B.1.351-challenged wildtype mice showed a milder infection than transgenic mice, confirmed by detectable viral loads in oropharyngeal swabs and tissues, lung pathology, immunohistochemistry and serology. *In silico* models supported these findings by demonstrating that the Spike mutations in B.1.351 resulted in increased affinity for both human and murine ACE2 receptors. Overall, this study highlights the plasticity of SARS-CoV-2 animal susceptibility landscape, which may contribute to viral persistence and expansion.

## Introduction

Severe Acute Respiratory Syndrome Coronavirus 2 (SARS-CoV-2) is a virus of zoonotic origin responsible for the Coronavirus Disease 2019 (COVID-19) that emerged by the end of 2019 in Wuhan, China (Holmes et al., 2021). From there, it rapidly spread across the world fuelling an unprecedented pandemic. The year 2020 ended with a growing awareness on SARS-CoV-2 variants that could potentially be more infectious and/or could evade immune responses elicited by previous infection with predecessor variants and/or vaccination (Trinité et al., 2021; Wang et al., 2021a).

To date, mass genotyping has prompted the identification of several variants of concern (VOC) (Mullen et al., 2020; Wang et al., 2021b): B.1.1.7/alpha —first described in the UK— showed higher transmissibility; B.1.351/beta, initially described in South Africa and P.1/gamma, isolated in Brazil, both demonstrated increased resistance to neutralizing antibodies elicited by other variants and/or vaccination (Dejnirattisai et al., 2021; Planas et al., 2021). More recently, the B.1.617.2/delta variant was identified in India and seems en route to rapidly supplant all other described SARS-CoV-2 variants (Parums, 2021). B.1.351’s greater resistance to neutralisation is especially concerning since it could considerably impact on vaccination programs (Shen et al., 2021). Furthermore, B.1.617.2 variant shows a combination of both higher transmissibility and resistance to neutralization (Li et al., 2021). Mutations in the spike (S) protein are responsible for these phenotypes by modifying neutralising epitopes and/or increasing the affinity for the human angiotensin converting enzyme 2 (hACE2) receptor (Gan et al., 2021). In addition to enhance the transmissibility between humans, these mutations can also alter the susceptibility of other host species of the virus, therefore broadening the animal reservoir (Conceicao et al., 2020; Liu et al., 2021; Piplani et al., 2021).

Also, animals are well-known coronavirus reservoirs (Menachery et al., 2017). While the intermediate host species of SARS-CoV-2 remains elusive, transmission from humans to domestic, farm and zoo animals has been documented (Muñoz-Fontela et al., 2020; Oreshkova et al., 2020; Segalés et al., 2020). The landscape of susceptibility species is also governed by S protein mutations, which can modulate the affinity to animal ACE2 (Garry, 2021). Notably, it has already been reported that new variants such as B.1.351 can infect SARS-CoV-2 resistant species like mice (Montagutelli et al., 2021). However, a complete *in vivo* susceptibility analysis has not yet been carried out.

The resistance of wildtype (WT) mice to SARS-CoV-2 infection made transgenic hACE2 mice one of the main experimental models for the *in vivo* study of novel vaccines and treatments (Bao et al., 2020; Jiang et al., 2020). The K18-hACE2 strain, a model used for SARS-CoV studies, is susceptible to SARS-CoV-2 infection and develops a severe disease. In these mice the hACE2 transgene is driven by the cytokeratin-18 (K18) gene promoter and hence it is expressed in many tissues, like lungs, intestines, and brain (Zheng et al., 2021). Widespread hACE2 expression drives a rapid progression of the disease, including massive brain infection, that does not recapitulate COVID-19 progression in humans (Winkler et al., 2020). Some groups, pursuing more physiological models, have attempted to mutate SARS-CoV-2 S protein to increase its tropism for mice (Gu et al., 2020; Sun et al., 2020). Alternatively, WT mice susceptibility to new SARS-CoV-2 variants, like B.1.351, may turn into an opportunity to develop a mouse model that better mimics the disease course, but may also represent a new viral reservoir.

The objective of this work was to compare the infectivity of B.1 and B.1.351 SARS-CoV-2 variants in both hACE2 transgenic and WT mice and characterise the progression of infection and pathological outcome. We provide evidence that B.1.351 displays a faster disease progression in hACE2 transgenic mice, while expanding tropism to WT mice. These results were consistent with molecular modelling suggesting an increased affinity of the B.1.351 associated S protein to both human and murine ACE2 and may have implications for experimental models of infection and, importantly, to SARS-CoV-2 ecology.

## Material & methods

### Biosafety Approval & Virus Isolation

The biologic biosafety committee of Germans Trias i Pujol Research Institute (IGTP) approved the execution of SARS-CoV-2 experiments at the BSL3 laboratory of the Centre for Bioimaging and Comparative Medicine (CMCiB, Badalona, Spain). The SARS-CoV-2 variants used in this study were isolated from nasopharyngeal swabs of hospitalised patients in Spain as described elsewhere (Perez-Zsolt et al., 2021; Rodon et al., 2021). Briefly, viruses were propagated in VeroE6 cells (CRL-1586; ATCC, Virginia, VA, USA) for two passages and recovered by supernatant collection. The sequences of the two SARS-CoV-2 variants tested here were deposited at the GISAID Repository (http://gisaid.org) with accession IDs EPI_ISL_510689 and EPI_ISL_1663571, respectively. EPI_ISL_510689 was the first SARS-CoV-2 virus isolated in Catalonia on March 2020 and, compared to the Wuhan/Hu-1/2019 strain, this isolate had the Spike mutations D614G, R682L, which are associated to the B.1 lineage. EPI_ISL_1663571 was isolated in February 2021 and showed the mutations associated with the B.1.351 or beta variant, and hence it will be referred as B.1.351. Viral stocks were titrated to use equivalent TCID50/mL on Vero E6 cells as described previously (Rodon et al., 2021).

### Animal Procedures and Study Design

All animal procedures were performed under the approval of the Committee on the Ethics of Animal Experimentation of the IGTP and the authorisation of *Generalitat de Catalunya* (code: 11222).

B6.Cg-Tg(K18-ACE2)2Prlmn/J (or K18-hACE2) hemizygous transgenic mice (034860; Jackson Immunoresearch, West Grove, PA, USA) were bred at CMCiB by pairing hemizygous males for Tg(K18-ACE2)2Prlmn (or K18-hACE2) with non-carrier B6.Cg females. The genotype of the offspring regarding Tg(K18-ACE2)2Prlmn was determined by qPCR at the IGTP’s Genomics Platform. Animals were kept in the BSL-3 facility during the whole experiment.

A total of 25 adult mice were used in this experiment (17 K18-hACE2 hemizygous mice and 8 non-transgenic WT mice from the same breed). All groups (4 to 7 animals) were sex balanced. Mice were anaesthetised with isoflurane (FDG9623; Baxter, Deerfield, IL, USA) and challenged with either B.1 or B.1.351 SARS-CoV-2 isolates, or PBS (uninfected control group). Infection was performed using 1000 TCID50 in 50 μL of PBS (25 μl/nostril). Mice fully recovered from the challenge and anaesthesia procedures.

After SARS-CoV-2 challenge, body weight and clinical signs were monitored daily. Two to three animals per group were euthanised 3 days post-infection (dpi) for viral load quantification. The rest of the animals were euthanised 7 dpi according to the experimental design (Supplementary Figure 1a), or earlier if they reached humane endpoint (body weight loss superior to 20% of the initial body weight and/or the display of moderate to severe neurological signs resulting from brain infection). Euthanasia was performed under deep isoflurane anaesthesia by whole blood extraction via cardiac puncture and was confirmed by cervical dislocation. Serum was recovered from whole blood following 10 min centrifugation at 4,000 x g. Lung, brain and nasal turbinate were collected for viral load determination and immunohistochemistry analysis.

### Quantification of humoral responses against SARS-CoV-2

The humoral response against SARS-CoV-2 was evaluated using an in-house ELISA, as previously reported (Brustolin et al., 2021). All samples were tested in the same plate in antigen coated and antigen-free wells for background subtraction. Briefly, half Nunc Maxisorp ELISA plates were coated with 50 μL of Spike, RBD or nucleocapsid protein (NP, SinoBiologicals, Beijing, China), all at 1 μg/mL in PBS and incubated overnight at 4°C. Then, plates were washed and blocked using PBS/1% of bovine serum albumin (BSA, Miltenyi biotech) for two hours at room temperature. After that, duplicates of each sample were added in the same plate to the wells coated with and without antigen. Samples were assayed at 1/200 (IgG determination) or 1/50 (IgM determination) dilution in blocking buffer overnight at 4°C. The next day, after washing, plates were incubated for 1 hour at room temperature with HRP conjugated-(Fab)2 Goat antimouse IgG (Fc specific) or Goat anti-mouse IgM (both 1/10000, Jackson Immunoresearch). Then, plates were washed and revealed with o-Phenylenediamine dihydrochloride (OPD, Sigma Aldrich) and stopped using 2N of H_2_SO_4_ (Sigma Aldrich). The signal was analyzed as the optical density (OD) at 492 nm with noise correction at 620 nm. The specific signal for each antigen was calculated as corrected optical density after subtracting the background signal obtained for each sample in antigen-free wells.

### SARS-CoV-2 PCR Detection and Viral Load Quantification

Viral load was determined in several tissues (lung, brain, oropharyngeal swab and nasal turbinate). Briefly, a piece of each tissue (100 mg approximately) was collected in 1.5 mL Sarstedt tubes (72,607; Sarstedt, Nümbrecth, Germany) containing 500 μL of DMEM medium (11995065; ThermoFisher Scientific) supplemented with 1% penicillin-streptomycin (10378016; ThermoFisher Scientific, Waltham, MA, USA). A 1.5 mm Tungsten bead (69997; QIAGEN, Hilden, Germany) was added to each tube and samples were homogenised twice at 25 Hz for 30 sec using a TissueLyser II (85300; QIAGEN) and centrifuged for 2 min at 2,000 x g. Supernatants were stored at −80 °C until use. RNA extraction was performed by using Viral RNA/Pathogen Nucleic Acid Isolation kit (A42352, ThermoFisher Scientific), optimized for a KingFisher instrument (5400610; ThermoFisher Scientific), following manufacturer’s instructions. PCR amplification was based on the 2019-Novel Coronavirus Real-Time RT-PCR Diagnostic Panel guidelines and protocol developed by the American Center for Disease Control and Prevention. Briefly, a 20 μL PCR reaction was set up containing 5 μL of RNA, 1.5 μL of N2 primers and probe (2019-nCov CDC EUA Kit, cat num 10006770, Integrated DNA Technologies, Coralville, IA, USA) and 10 μl of GoTaq 1-Step RT-qPCR (Promega, Madison, WI, USA). Thermal cycling was performed at 50°C for 15min for reverse transcription, followed by 95°C for 2 min and then 45 cycles of 95°C for 10 sec, 56°C for 15 sec and 72°C for 30 sec in the Applied Biosystems 7500 or QuantStudio5 Real-Time PCR instruments (ThermoFisher Scientific). For absolute quantification, a standard curve was built using 1/5 serial dilutions of a SARS-CoV2 plasmid (2019-nCoV_N_Positive Control, catalog number 10006625, 200 copies/μL, Integrated DNA Technologies) and run in parallel in all PCR determinations. The viral load of each sample was determined in triplicate and mean viral load (in copies/mL) was extrapolated from the standard curve and corrected by the corresponding dilution factor. Mouse *gapdh* gene expression was measured in duplicate for each sample using TaqMan^®^ gene expression assay (Mm99999915_g1; ThermoFisher Scientific) as amplification control. The 2^-ΔCT^ value was calculated for each sample.

### Histopathological and Immunohistochemical analyses

Formalin-fixed lung, nasal turbinate and brain samples were routinely processed for histopathology, and haematoxylin&eosin-stained slides were examined under optical microscope. A semi-quantitative approach based on the amount of inflammation (none, mild, moderate, or severe) was used to establish the damage caused by SARS-CoV-2 infection in mice, following a previous published scoring system (Vidal et al., submitted).

An immunohistochemistry (IHC) technique to detect SARS-CoV-2 nucleoprotein (NP) antigen using the rabbit monoclonal antibody (40143-R019, Sino Biological, Beijing, China) at a 1:15,000 dilution, was applied on nasal turbinate, lung and brain sections from all animals. The amount of viral antigen in tissues was semi-quantitatively scored (low, moderate and high amount, or lack of antigen detection) (Vidal et al., submitted).

### Molecular Modelling of ACE2 species with SARS-CoV-2 Spike

Since no experimental structure of murine ACE2 (mACE2) in complex with the spike protein of SARS-CoV-2 was available, we ran MODELLER v10.1 (Šali and Blundell, 1993) to generate homology models. As a template, the crystallographic structure of hACE2 in complex with the spike RBD of the initial virus variant (PDB id 6M0J) was used. We created a total number of 10 models (identified as mACE2-WT RBD). In a second step, using the MODELLER models as input, we ran FoldX v5 (Schymkowitz et al., 2005; Delgado et al., 2019) to model the mutations associated with the B.1.351 variant of the virus, obtaining 10 additional models (named mACE2-B.1.351 RBD) (Supplementary Fig. 3). All FoldX parameters had default values except parameter vdwDesign that we set to zero. We evaluated each model with FoldX and pyDock (Cheng et al., 2007) which have achieved good performance predicting the impact of mutations in protein-protein complexes (Amengual-Rigo et al., 2021). As a control, we repeated the procedure described above to generate models of hACE2 - in complex with the RBD of the WT virus (hACE2 – WT RBD) and the B.1.351 variant (hACE2 – B.1.351 RBD). As before, we ended up with 10 different models per complex type. We visualized and produced the pictures of the models with UCSF Chimera (Pettersen et al., 2004).

## Results & Discussion

### B.1.351 develops a faster and more severe form of the disease in K18-hACE2 mice than the B.1 VOC

K18-hACE2 transgenic mice were experimentally inoculated with SARS-CoV-2 variant B.1 (n=6) or B.1.351 variant (n=7). An uninfected control group was established by challenging four K18-hACE2 animals with PBS-only (n=4). At 3 dpi, two animals challenged with the B.1 and three animals challenged with B.1.351 were euthanised for early viral load quantification in tissues, while four animals per group were followed-up until day 7 or until reaching humane endpoint (Supplementary Figure 1a).

B.1-challenged K18-hACE2 animals that were followed up until 7 dpi (n=4) started losing body weight on 5 dpi (Figure 1a) and three of them met humane endpoint criteria on day 6 (n=1) and 7 (n=2) post-infection (Supplementary Figure 1b,c). In comparison, B.1.351-challenged K18-hACE2 mice started losing body weight as early as 3 dpi (Figure 1a), displaying an accelerated disease progression leading to euthanasia on days 5 (n=3) and 6 (n=1) post-infection (Supplementary Figure 1b,c).

**Figure 1.**
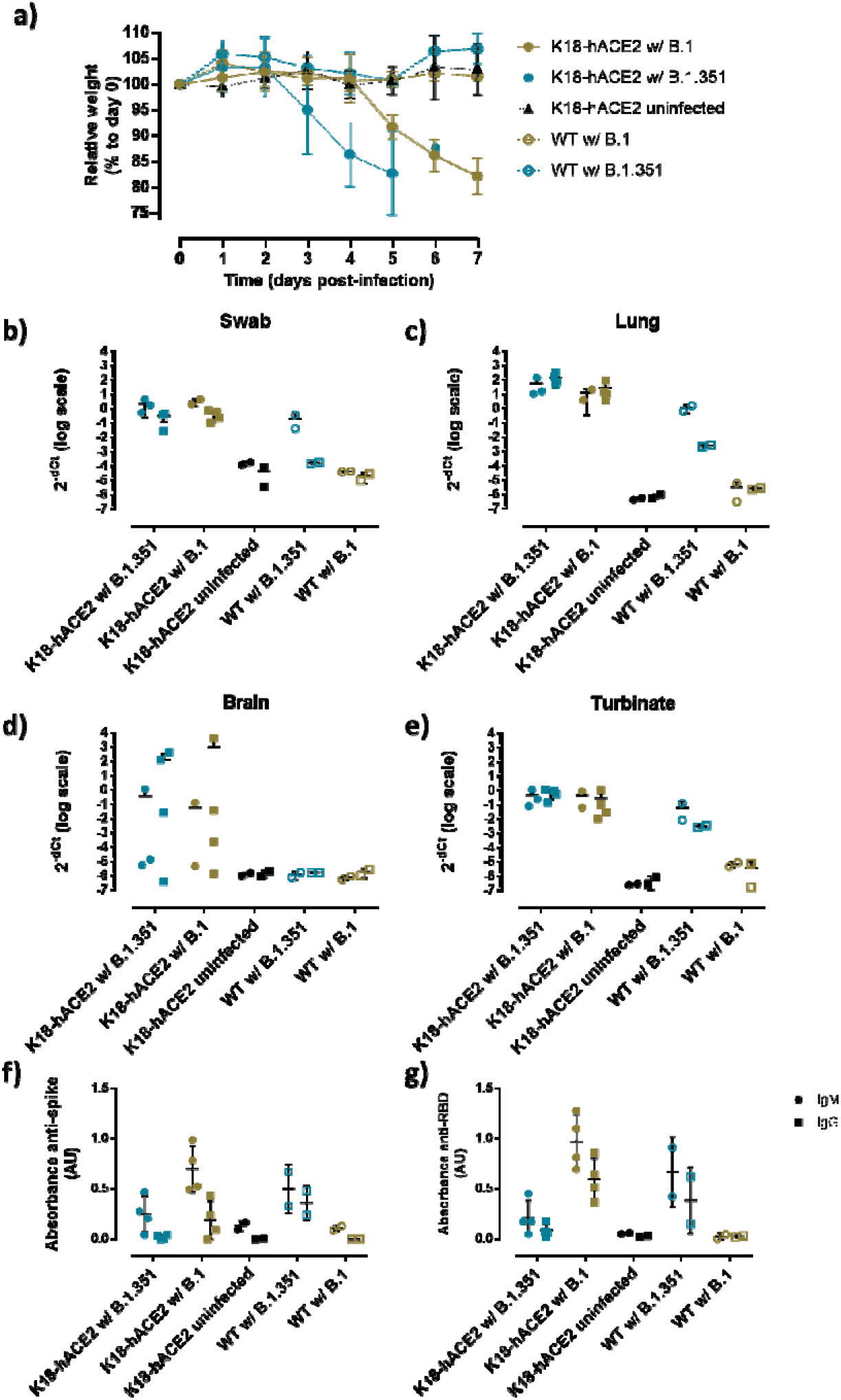
Disease progression and monitorisation of K18-hACE2 and wildtype (WT) mice inoculated with SARS-CoV-2 B.1 and B.1.351 variants. a) Relative body weight follow-up of K18-hACE2 mice (solid symbols) and WT mice (empty symbols) infected with B.1 (gold), B.1.351 (blue) or uninfected (black). b-e) SARS-CoV-2 viral loads represented as 2^-ΔCt^ on different tissues: b) swab; c) lung; d) brain; e) nasal turbinate. Round shapes represent animals euthanised on day 3, while squares represent viral loads on animals euthanised at endpoint (days 5-7). f-g) Serological analysis of IgG and IgM induced in infected and uninfected animals at the day of euthanasia (days 5-7) against: f) S protein (sera dil. 1/200), and g) RBD (sera dil. 1/50).

Viral load in lung, brain and nasal turbinate revealed widespread infection in K18-hACE2 transgenic mice, with no relevant differences detected between B.1- and B.1.351-challenged animals (Figure 1b-e). All tissues contained remarkably high viral loads, except for the brain, where SARS-CoV-2 viral loads were more heterogeneous regardless of the variant (Figure 1d).

Analysis of the humoral response against SARS-CoV-2 Spike protein and the RBD measured at endpoint showed detectable levels of IgM in both B.1 and B.1.351-infected K18-hACE2 mice, although IgM titres were lower in the latter (Figure 1f,g). In contrast, only B.1-infected animals developed IgGs at the time of euthanasia. Both observations could be attributed to the faster disease progression in B.1.351-infected transgenic mice, which would not allow for development of a full-fledged immune response before euthanasia.

The histopathological analysis showed no evident inflammatory lesions in the nasal turbinate of any of the inoculated mice, irrespective of the challenging isolate (Supplementary Figure 2). The mean viral antigen score measured by IHC on 3 dpi was slightly higher for K18-hACE2 mice inoculated with the B.1.351 (two animals with a score of 1 and one animal with a score of 2) compared to those challenged with B. 1 (two animals with a score of 1). By 7 dpi, all four B.1.351-challenged mice and three out of four B.1-inoculated animals had an IHC score of 1.

Lungs of K18-hACE2 inoculated mice showed mild to moderate interstitial pneumonia on 3 dpi and at endpoint, with one animal challenged with B.1.351 showing a severe lesion at endpoint (Figure 2a). IHC results in lung showed moderate to high amounts of viral antigen on 3 dpi and endpoint in both groups of K18-hACE2 challenged mice (Figure 2b,c).

**Figure 2.**
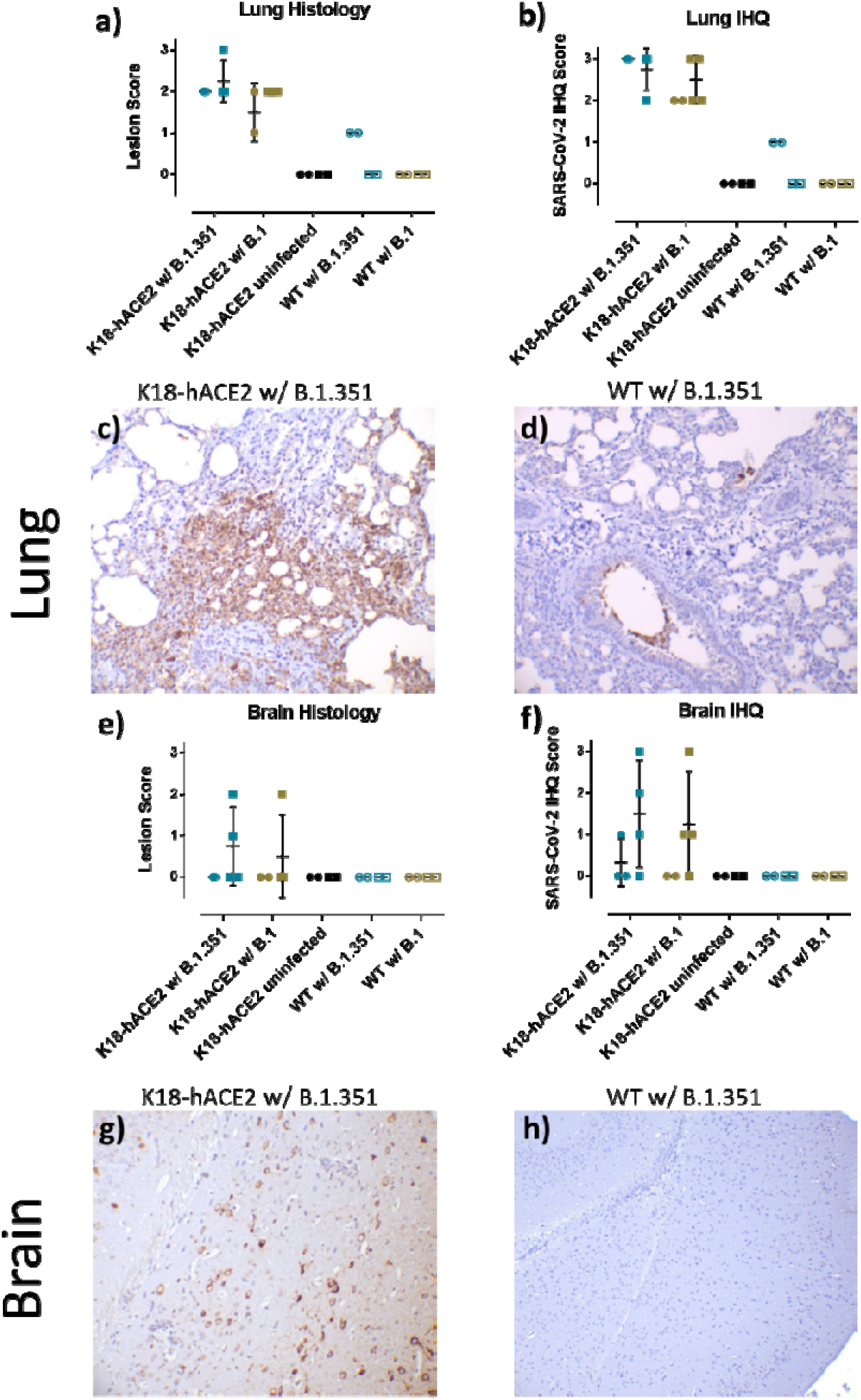
Histopathological and immunohistochemical analyses of K18-hACE2+ and K18-hACE2-mice inoculated with SARS-CoV-2 variants B.1 and B.1.351. a) Lung histopathological and b) immunohistochemical scores of individual animals of each experimental group. c) High amount of SARS-CoV-2 antigen found in lung parenchyma displaying interstitial pneumonia in a hACE2+ mouse inoculated with the B.1.351 isolate. d) Low amount of SARS-CoV-2 antigen detected in few epithelial respiratory cells as well as in sloughed cells into bronchus lumen. e) Brain histopathological and f) immunohistochemical scores of individual animals of each experimental group. g) High amount of viral antigen within neurons of the brain of the hACE2+ mice inoculated with B.1.351 and showing moderate non-suppurative meningo-encephalitis. h) Lack of viral immunolabelling in the brain of an hACE2-mouse inoculated with the B.1.351 variant. Horizontal short bars in figures a, b, e and f represents the average of the corresponding group. All immunohistochemical slides were Haematoxylin counterstained.

The brain of two K18-hACE2 mice infected with B.1.351 and one K18-hACE2 mice infected with B.1 showed mild to moderate non-suppurative meningoencephalitis at endpoint (Figure 2e). No brain lesions were observed in any other animal from the study. IHC results in brain on 3 dpi were restricted to one single K18-hACE2 mouse challenged with B.1.351 (Figure 2f), with a low amounts of viral antigen in neurons. In contrast, three K18-hACE2 animals from each of the two SARS-CoV-2 inoculated groups had low to high amount of viral antigen at endpoint (Figure 2f,g).

### B.1.351 can infect wild type mice promoting a mild form of disease

In parallel to the challenge experiment described above, K18-hACE2 negative mice from the same breed WT were also challenged with the B.1 (n=4) or B.1.351 (n=4) variants. The results showed no body weight alterations until 7 dpi in none of the WT infected animals (Figure 1a), similar to uninfected mice. Viral load quantification demonstrated that B.1.351 was able to infect the lungs and the nasal turbinate of WT mice (Figure 1c,e). Viral load was also detectable in oropharyngeal swabs at 3 dpi (Figure 1b). The B.1 variant was not detected by qRT-PCR in any analysed tissue at any timepoint (Figure 1c-e). Moreover, anti-Spike and anti-RBD IgM and IgG antibodies were observed only in B.1.351-infected WT mice but not in those challenged with the B.1 variant (Figure 1f,g), further supporting the fact that B.1 could not effectively infect WT mice, while B.1.351 did.

Two relevant aspects of B.1.351 SARS-CoV-2 infection in WT mice were that virus could not be detected in the brain (Figure 1d), and that viral loads from the swab, lung and nasal turbinate of animals euthanised 3 dpi were higher than viral loads in the samples recovered at endpoint (Figure 1b,c,e). This could explain why, despite getting infected, WT animals did not display severe signs as hACE2-expressing mice did. Reduction of viral loads in WT mouse tissues may reflect a rapid and effective viral clearance by the immune response (Figure 1f,g), or the incapacity of the virus to sustain multiple cycles of infection.

Similar to what was observed in hACE2-expressing mice, no inflammatory lesions were observed in nasal turbinate of both B-1 and B.1.351 challenged WT mice (Supplementary Figure 2). Viral antigen was only detected by IHC (score 1) in one out of two animals challenged with B.1.351 at 3 dpi. The rest of animals (one from 3 dpi and two from 7 dpi) were negative by IHC. Interestingly, WT animals inoculated with B.1.351 showed mild interstitial pneumonia by 3 dpi. However, no lung lesions were observed in WT mice inoculated with B.1 at any time post-challenge or in negative control ones (Figure 2a). WT mice inoculated with the B.1.351 isolate showed low amount of viral antigen labelling in lung tissue at 3 dpi (Figure 2b,d), while it was not detected by IHC in WT mice inoculated with B.1.351 at 7 dpi, uninfected controls and WT mice inoculated with B.1. Negative control and WT mice inoculated with both variants had no labelling in brain at any time post-inoculation (Figure 2f,h).

### Molecular modelling of ACE2 from mice with SARS-CoV-2 Spike

To estimate the effect of the mutations associated with the B.1.351 variant on the binding affinity to ACE2, the protein-protein interaction models were evaluated with two levels of theory: FoldX and pyDock. Both models yielded significantly lower energies (i.e., higher binding affinity predictions) for the B.1.351 variant spike RBD in complex with both human and murine ACE2 (Wilcoxon paired test *p*=0.01) (Figure 3). These results suggest that the effect of B.1.351 mutations could improve binding to human and murine ACE2.

**Figure 3.**
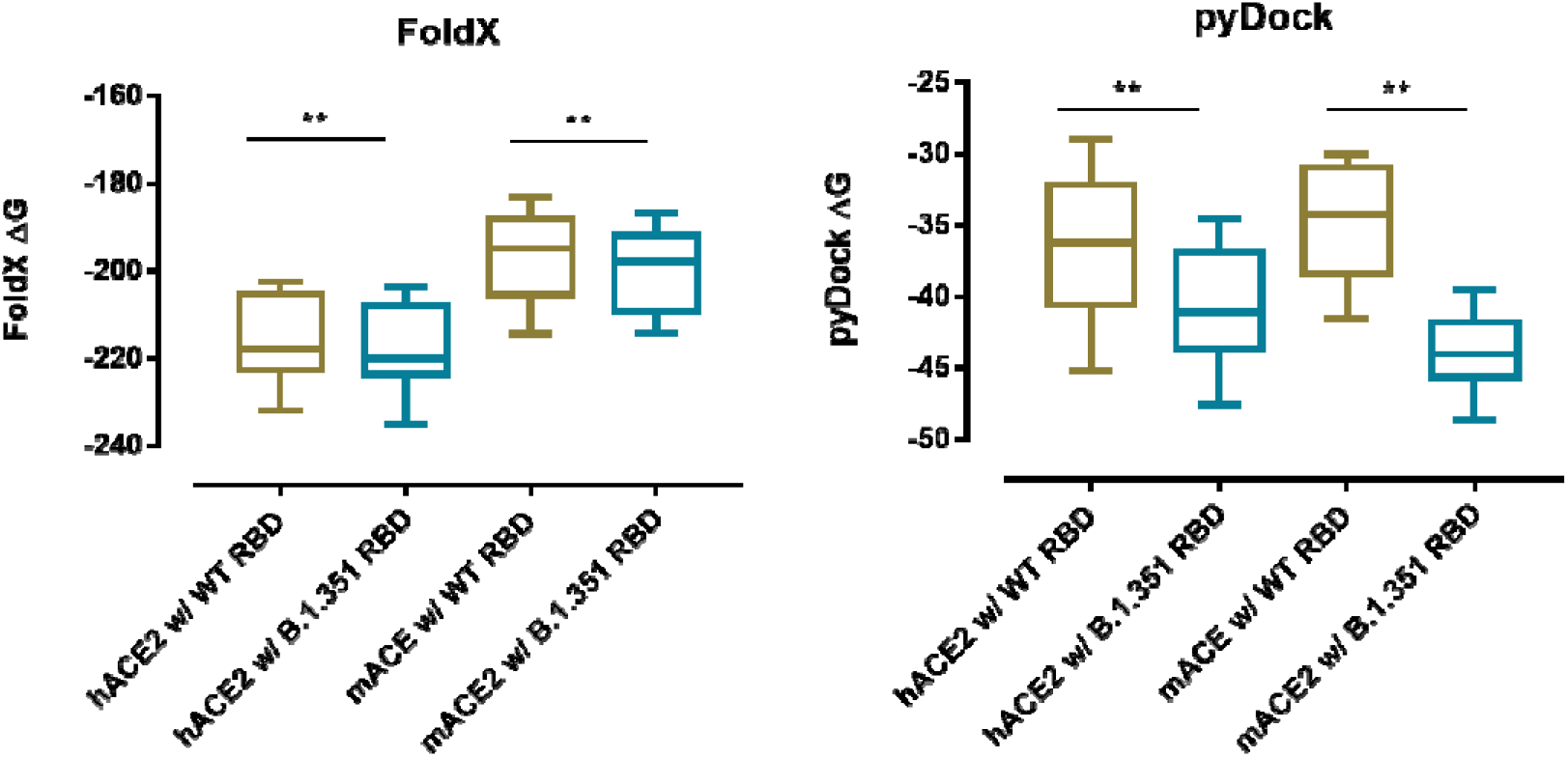
FoldX (left) and pyDock (right) energies of the models of hACE2 and mACE2 in complex with the initial (Wuh) and B.1.351 variants of the SARS-CoV-2 spike RBD. Complexes with lower energies are predicted as having higher binding affinities by both methods. FoldX and pyDock energies are significantly lower for the beta variant spike RBD in complex with ACE2 in human and mice. Significant differences were computed with a Wilcoxon paired test.

A detailed analysis of the modelled structures provided a mechanistic description of the mutation effects at the molecular level. The exploration of the models suggested that the mutations did not produce significant structural changes (Supplementary Figure 4). Mutation K417N caused the loss of a salt bridge between Lys417 and hACE2 Asp30, reducing the affinity to the human receptor. Interestingly, in mice, no salt bridge is formed between Lys417 and mACE2 Asn30, and hence binding affinity is not negatively affected by the mutation (Supplementary Figure 4a-d). A similar case occurs with mutation E484K, which induces the loss of a salt bridge between Glu484 and Lys31 in hACE2, but not in mACE2, where this salt bridge cannot be formed between Glu484 and mACE2 Asn31 (Supplementary Figure 4e-h). Mutation N501Y increases the number of hydrophobic interactions between Tyr501 and ACE2 Tyr41 both in human and mice. Additionally, Tyr501 can establish a cation-pi interaction with hACE2 Lys353 in human and a pi-pi interaction with mACE2 His353 in mice (Supplementary Figure 4i-l). In summary, mutation N501Y should increase the affinity in both species. This effect would be reduced by the loss of salt bridges caused by mutations K417N and E484K in human, but not in mice.

Altogether, molecular analyses suggest that multiple mutations within the B.1.351 SARS-CoV-2 Spike enhanced the affinity for mACE2, which might explain why WT mice could get infected by this variant.

### Conclusions

This study shows that the SARS-CoV-2 B.1.351 variant can establish infection in WT mice, in line with previous findings (Montagutelli et al., 2021). Compared to hACE2-expressing transgenic mice that developed a severe clinical course and pathology upon SARS-CoV-2 infection, WT mice displayed a milder phenotype. This milder outcome could be comparable to SARS-CoV-2 infection in other animal models, in which lung replication and viral pneumonia is detected, but eventually resolved by the action of the immune response (Brustolin et al., 2021), and could open the door to develop better SARS-CoV-2 models than transgenic hACE2 mice. This, however, also raises concerns on how new variants with increased affinity to ACE2 could expand their host tropism to animal species that were non-susceptible to earlier SARS-CoV-2 variants, potentially generating new zoonotic reservoirs and increasing the risk of new spillover and spread of new variants in humans.

Further studies will prove whether, in a natural setting, increased affinity to mACE2 by B.1.351 mutations can result in an effective SARS-CoV-2 infection, or whether the mild symptoms and viral control in WT mice caused by B.1.351 were just due to the virus’ inability to sustain infection. Still, this study confirms the need to monitor SARS-CoV-2 VOC beyond its interaction with humans (transmissibility, pathogenicity and immune evasion) and from a “One Health” perspective.

## Acknowledgements

The authors would like to acknowledge J. Díaz from the CMCiB for his constant help at the BSL3 facility and Y. Rosales and R. Ampudia from the CMCiB for their unvaluable help with the K18-hACE2 colony. We also thank M. Parera from IrsiCaixa for her outstanding help in sequencing viral variants.

## Supplementary figures

**Supplementary Figure 1.**
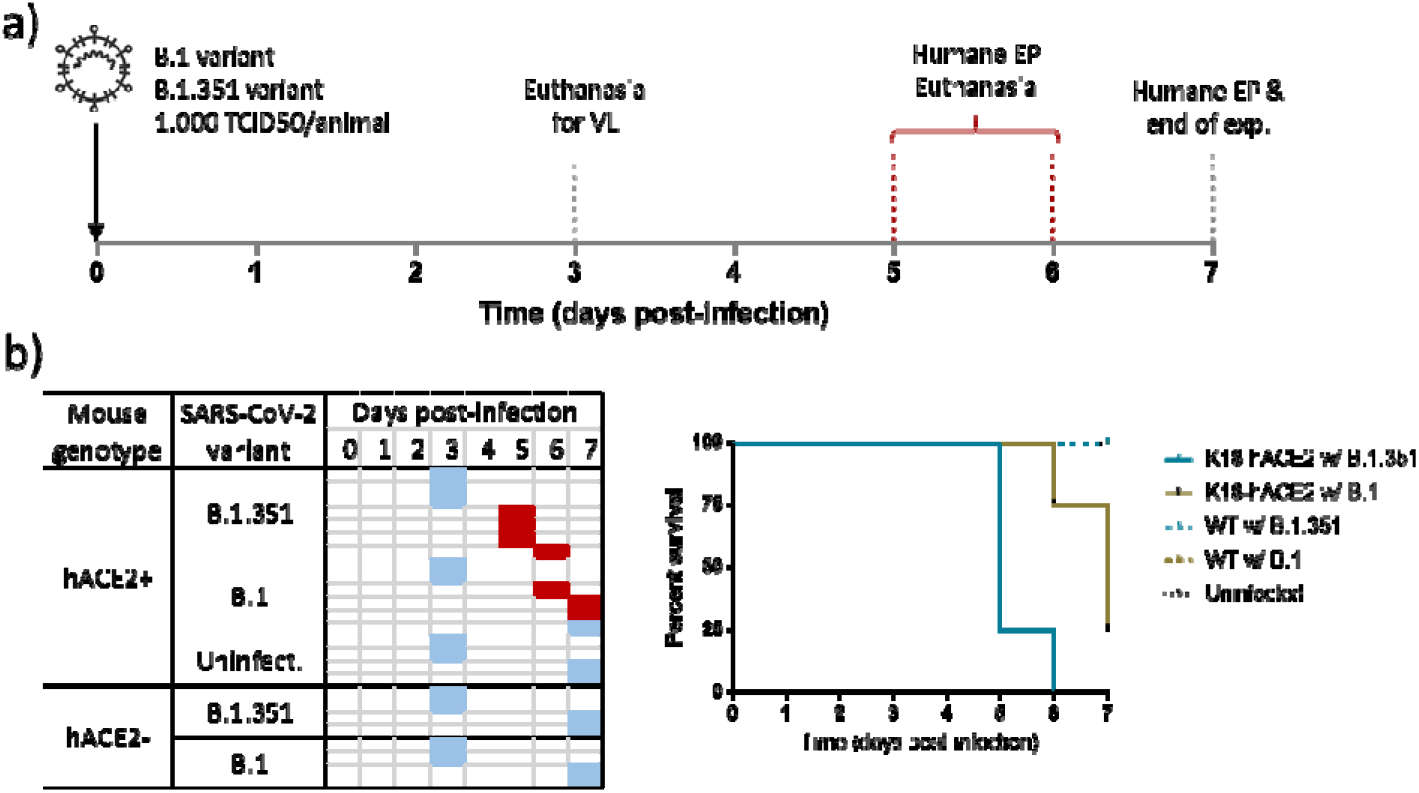
Experimental design and euthanasia timepoints of K18-hACE2 and WT mice inoculated with SARS-CoV-2 variants B.1 and B.1.351. a) Schedule from day 0 to day 7. b) Euthanasia schedule for each animal; blue represents a scheduled euthanasia according to the protocol, while red represents euthanasia upon reaching humane endpoint criteria. c) Survival curve of K18-hACE2 animals infected with B.1 (gold) or B.1.351 (blue).

**Supplementary Figure 2.**
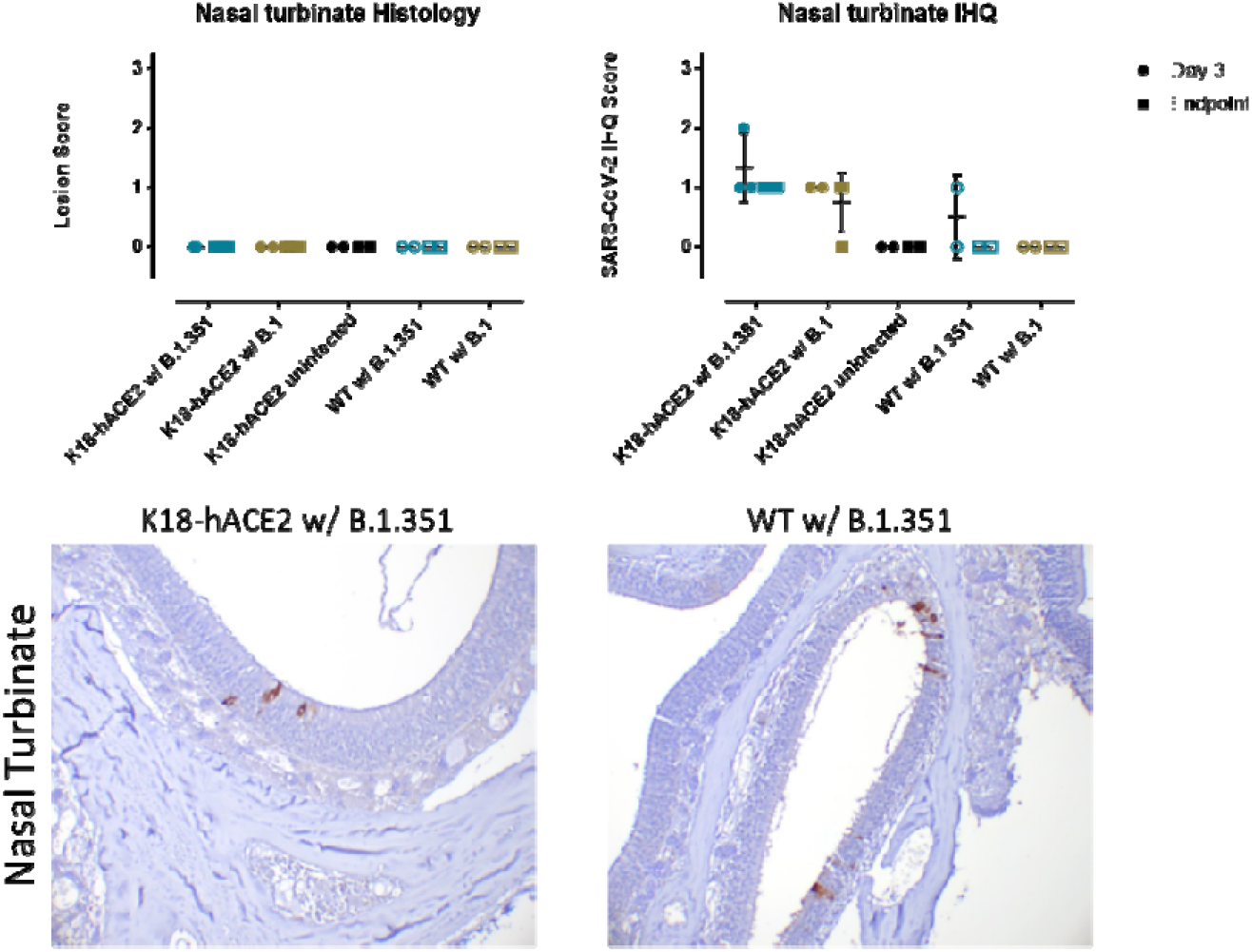
Lesions and replication in the nasal turbinate. K18-hACE2 (solid symbols) and WT mice (empty symbols) infected with SARS-CoV-2 variants B.1 (gold) and B.1.351 (blue). a) Histopathological and b) immunohistochemical scores of individual animals of each experimental group. c) Representative figure of the nasal turbinate of a K18-hACE2 mouse infected with B.1.351. d) Representative figure of the nasal turbinate of a WT mouse infected with B.1.351.

**Supplementary Figure 3.**
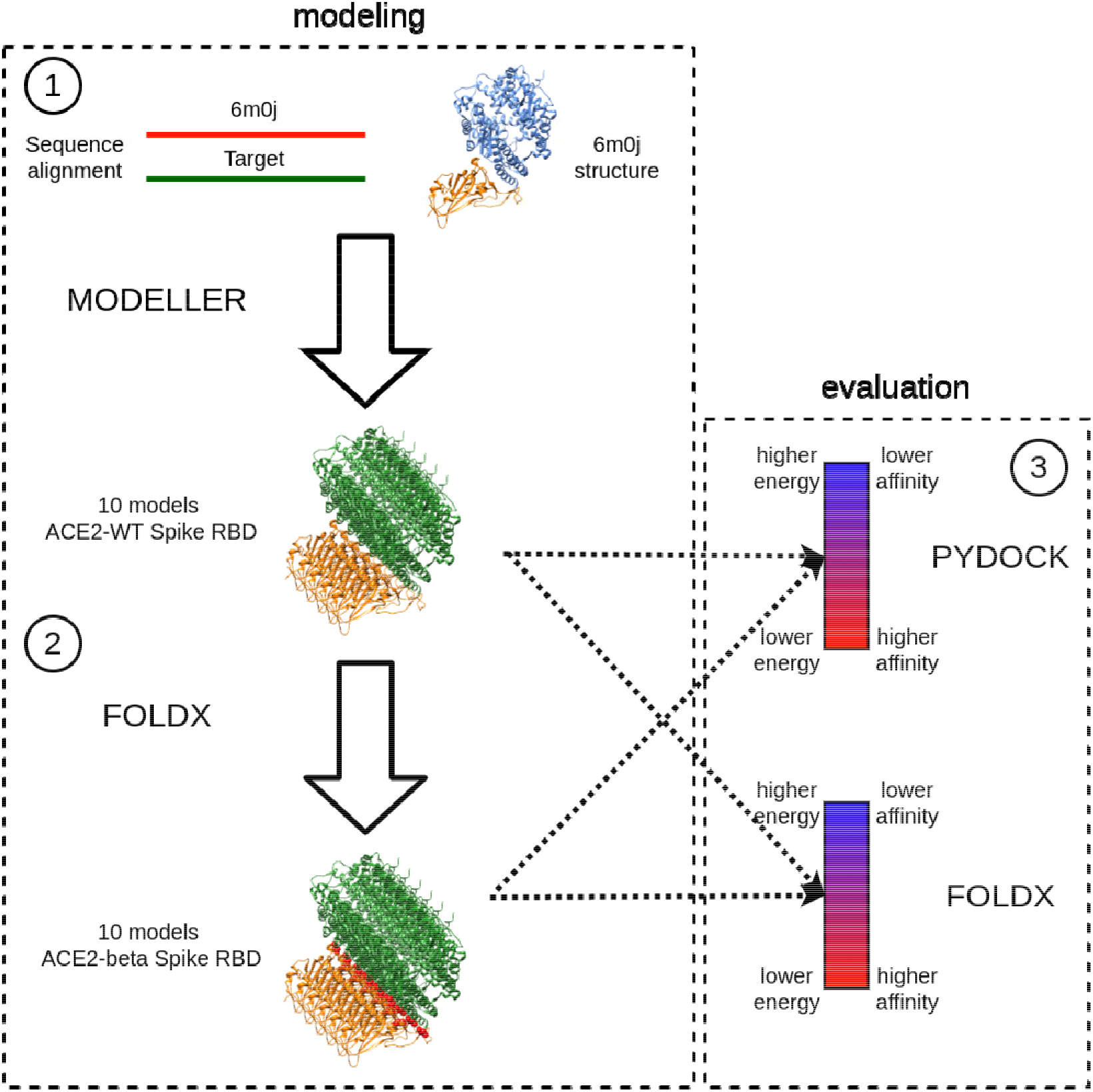
Protocol diagram. 1) Ten models of ACE2 in complex with the spike RBD of the initial WT virus variant were created with MODELLER. The structure with PDB id 6M0J was used as a template. 2) Mutations associated with the B.1.351 variant at the spike RBD were modelled with FoldX using MODELLER models as input. 3) All models were evaluated with FoldX and pyDock to obtain an estimation of the binding affinity changes due to the mutations. We ran the protocol twice: The first time to generate mACE2-RBD complexes, the second time to produce hACE2-RBD complexes to use as a control.

**Supplementary Figure 4.**
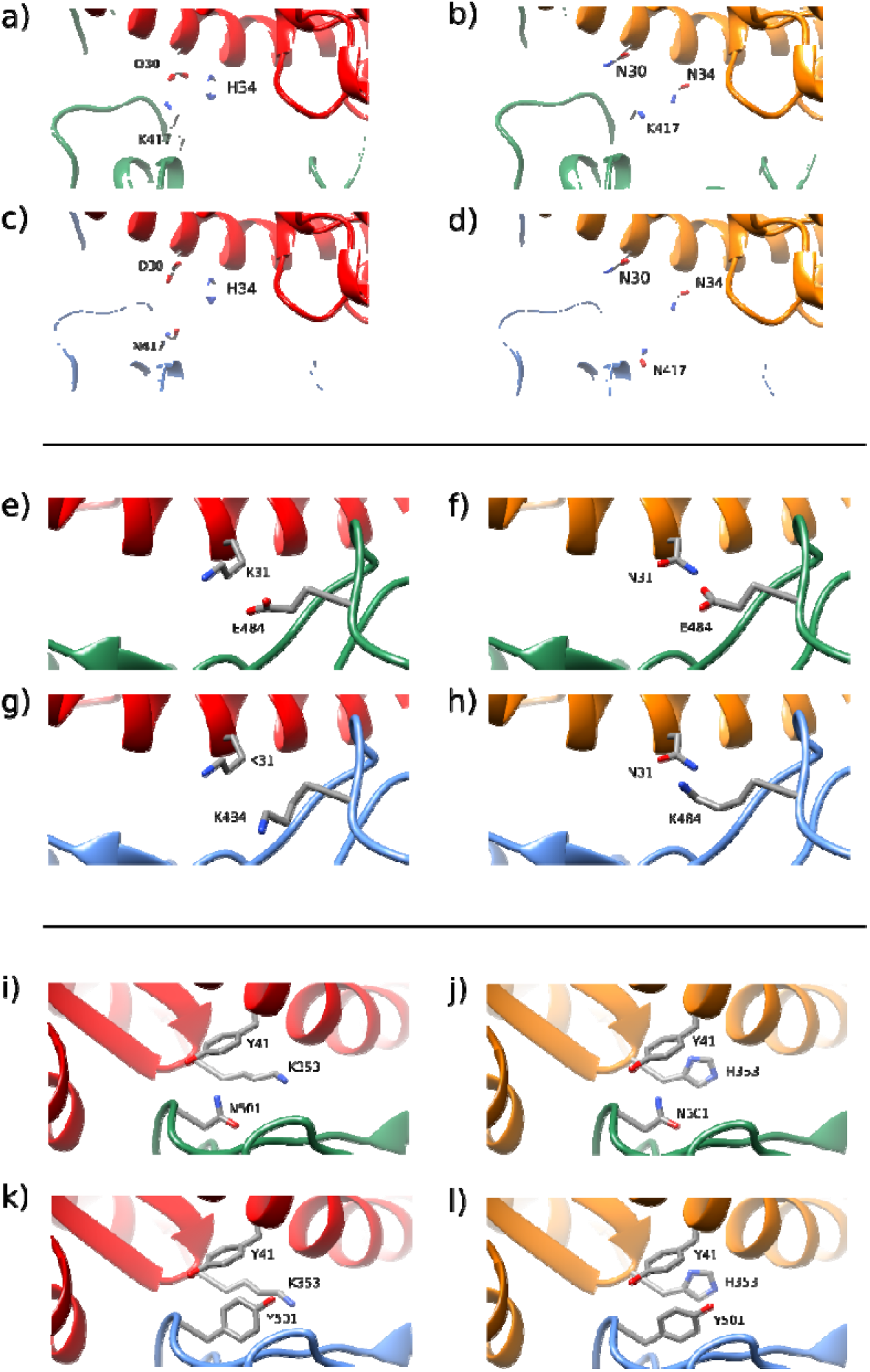
Structural details in models with hACE2 (red), mACE2 (orange), WT Spike (green) and B.1.351 Spike (blue). In human, mutation K417N causes the loss of a salt bridge between Lys417 and hACE2 Asp30, diminishing the binding affinity of the complex (a, c). In mice, no salt bridge is formed between WT RBD Lys417 and mACE2 Asn30, and affinity does not decrease by its loss (b, d). Similarly, mutation E484K induces the loss of a salt bridge between Glu484 and Lys31 in hACE2 (e, g), but not in mACE2, where no salt bridge is established between WT RBD Glu484 and mACE2 Asn31 (f, h). Mutation N501Y increases the number of hydrophobic interactions between Tyr501 and ACE2 Tyr41 in human and mice (i-l). Tyr501 can have a cation-pi interaction with hACE2 Lys353 in human (k), and a pi-pi interaction with mACE2 His353 in mice (l). The analysis suggests that the increase in affinity due to mutation N501Y is partially reduced by the loss of salt bridges in human, but not in mice.

